# Spatial Transcriptomics Reveals Spatially Diverse Cancer-Associated Fibroblast in Lung Squamous Cell Carcinoma Linked to Tumor Progression

**DOI:** 10.1101/2024.05.16.594592

**Authors:** Hongyoon Choi, Kwon Joong Na, Yeonjae Jung, Myunghyun Lim, Dongjoo Lee, Jae Eun Lee, Hyung-Jun Im, Daeseung Lee, Jaemoon Koh, Young Tae Kim

## Abstract

While cancer-associated fibroblasts (CAFs) are crucial in influencing tumor growth and immune responses in lung cancer, we still lack a comprehensive understanding of their spatial organization associated with tumor progression and clinical outcomes. This gap highlights the need to elucidate how the intricate spatial arrangement of CAFs affects their interactions within the tumor microenvironment, ultimately shaping cancer progression and patient prognosis. Here, we unveil the spatial diversity of CAFs in lung squamous cell carcinoma (LUSC), a prevalent and aggressive lung cancer type, elucidating their impact on tumor progression and patient outcomes using spatial transcriptomics (ST). Image-based ST data from 33 LUSC patients demonstrated a significant association of spatial interactions of tumor epithelium and CAFs with tumor size and metabolic activity measured by [^18^F]fluorodeoxyglucose PET. Furthermore, the proximity of fibroblasts to tumor epithelial cells was linked to recurrence-free survival in LUSC patients. By characterizing CAFs based on their spatial relationship, we identified distinct molecular signatures related to spatially distinct fibroblast subpopulations. In addition, barcode-based ST data from 8 LUSC patients revealed spatially overlapping fibroblast regions characterized by upregulated glycolysis pathways. Our study underscores the importance of the complex spatial dynamics of the tumor microenvironment revealed by ST and its implications for patient outcomes in LUSC.

## Main

Lung squamous cell carcinoma (LUSC) represents a particularly aggressive non-small cell lung cancer subtype, where the tumor microenvironment (TME) plays a critical role in its progression and response to therapy^1^. Within this microenvironment, cancer-associated fibroblasts (CAFs) are crucial contributors, significantly impacting tumor growth, immune evasion, and therapeutic resistance^2,3^. Recent single-cell RNA sequencing (scRNA-seq) studies have unveiled a remarkable diversity in CAF subpopulations within the microenvironment^4-7^. Notably, these CAFs exhibit a range of functions from extracellular matrix (ECM) remodeling and modulation of immune responses to direct effects on cancer cell proliferation and metastasis^4,6^. The interaction between LUSC cells and CAFs is intricate, involving various signaling pathways and cellular interactions that ultimately influence tumor behavior and patient outcomes^8-12^. Understanding the role of CAFs is especially crucial in the context of lung cancer, where the efficacy of immuno-oncology treatment hinges on the nuanced interplay of various cellular constituents within the tumor microenvironment^13-15^. However, the spatial organization and functional dynamics of CAFs within this setting remain largely unknown territories, yet they hold the key to unlocking novel therapeutic strategies and improving patient outcomes.

Spatial transcriptomics (ST), a groundbreaking approach that integrates spatial contexts of disease tissues with genomic data, offering a comprehensive view of the tumor landscape at a high resolution^16,17^. ST can be broadly categorized into two types based on the underlying techniques: barcode (or NGS)-based and image-based ST^18^. Barcode-based ST utilizes next-generation sequencing to analyze the entire genome, although it offers limited spatial resolution^17,19-21^. In contrast, image-based ST employs multiplex detection of transcripts using pre-designed gene panels, which provides high spatial resolution despite its restricted gene coverage^22-24^. By utilizing image-based and barcode-based ST techniques, researchers can now explore the spatial and molecular intricacies of disease tissue, including lung cancer. This method transcends the limitations of single-cell analyses, which, while informative, fail to capture the critical spatial relationships and patterns essential for understanding tumor behavior^25^. The application of ST in lung cancer research, particularly in LUSC, promises to shed light on the spatial heterogeneity of tumor microenvironment cells including CAFs and their interactive networks within the TME.

The purpose of this study is to dissect the spatial relationships of various cell types and molecular features in tumor microenvironment in LUSC. We employed ST and analyzed with clinical and imaging features of LUSC to unravel the complex spatial relationship which is highly associated with tumor progression and influence clinical outcomes. Our investigation into 33 LUSC patients reveals significant correlations between the spatial positioning of CAFs and key tumor characteristics, including tumor size and metabolic activity, as well as implications for prognosis. More specifically, by investigating the spatial relationship of cell types and molecular features of the tumor microenvironment from patients-derived ST data, we aimed to unravel the complex mechanisms driving LUSC progression. Our approach of ST analyses with clinical features could eventually offer a detailed spatial map of tumor epithelium-CAFs crosstalk and highlighting potential biomarkers. The suggested comprehensive approach to understanding the TME in spatial context not only advances our knowledge of LUSC pathophysiology but also paves the way for investigating a tumor ecosystem in the evolving field of cancer research.

## Results

### Image-based ST revealed spatial organization of spatial cell types in TME of LUSC

We embarked on a comprehensive analysis of the TME in LUSC by prospectively collecting surgical tissue samples from thirty-three patients. The demographics and clinicopathological features of these individuals are detailed in **Supplementary Table 1**. We constructed tissue microarrays (TMAs) using 2 mm cores from these samples to conduct image-based ST. Utilizing the multiplexed error-robust fluorescence in situ hybridization (MERFISH) technology, which incorporates a 500-gene panel designed to delineate TME cell types, we analyzed these TMAs to generate a high-resolution cellular atlas of the LUSC TME (as shown in **Figure 1a**). Each TMA core provided a distinct sample from the LUSC cases, allowing us to observe the expression patterns of critical marker genes within the microenvironment (**Figure 1a**).

**Figure 1.**
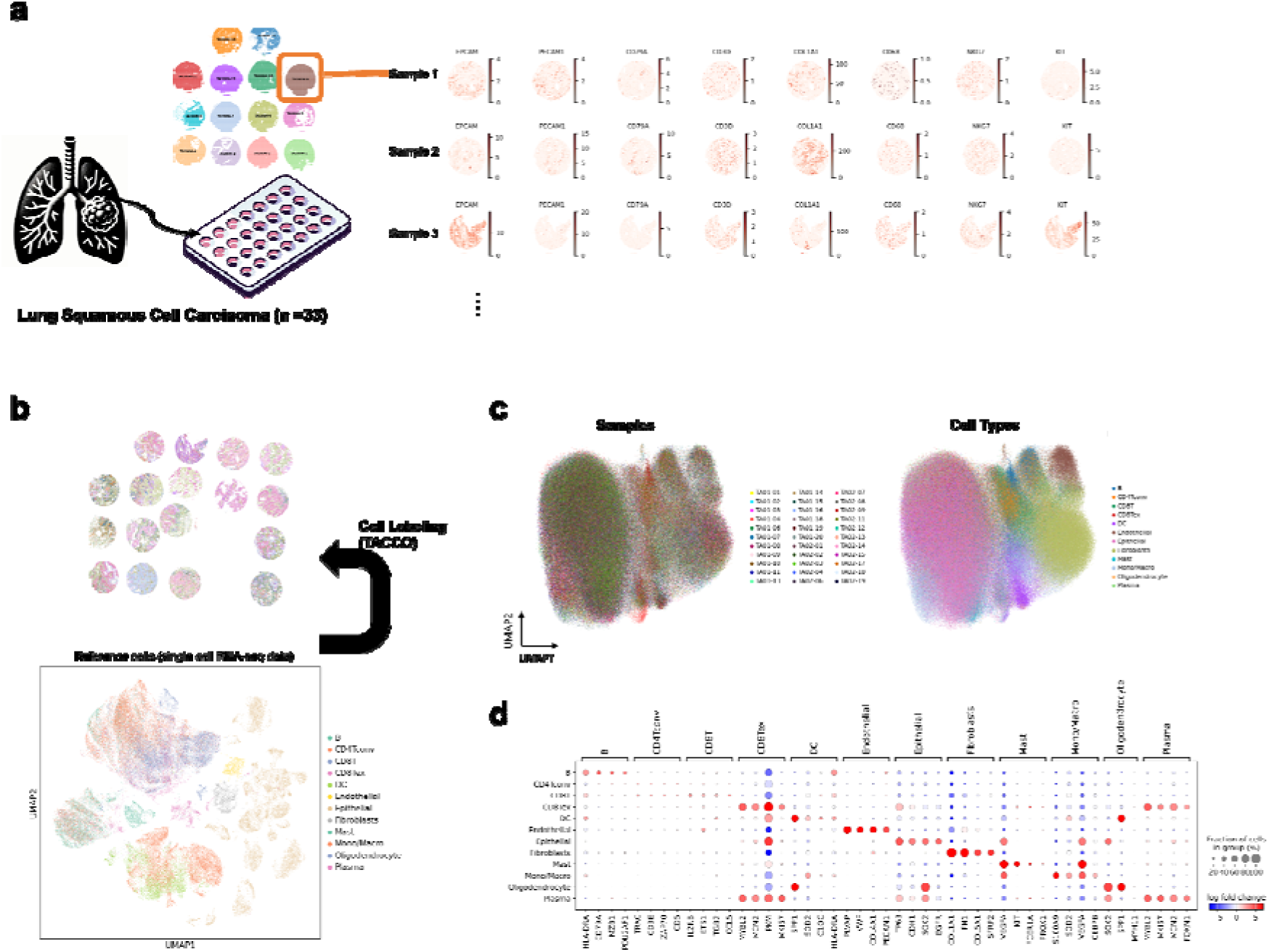
MERFISH for lung squamous cell carcinoma (LUSC) and mapping cell types in the tumor microenvironment (TME) (a) Tissue microarrays (TMAs) containing 2 mm cores from 33 LUSC surgical samples, analyzed using MERFISH. Custom designed 500 gene sets for analyzing spatial transcriptomic profiles were used to show the expression patterns of critical TME marker genes across different samples. (b) Application of the TACCO algorithm to assign cell types based on a reference single cell RNA-seq (scRNAseq) dataset. (c) UMAP plots depict different samples (*left*) and cell types (*right*), each color-coded to represent a distinct sample or cell type. (d) Differential gene expression analysis identifying key markers for each cell type in the MERFISH data. A dot plot visualizes the expression of these markers, further confirming cell type identifications based on the MERFISH gene expression data.

To assign cell types to each spatial location within our samples, we employed the transfer of annotations to cells and their combinations (TACCO) algorithm, an annotation transfer method based on single-cell RNA sequencing (scRNA-seq) data^26^. The reference dataset for this algorithm was sourced from a public database containing information on non-small cell lung cancer (GSE131907)^7^ (**Figure 1b**). The TACCO algorithm generated the probability of cell types for each cell in ST data, which encompassed 465,707 cells tagged with data on 500 gene expressions. Each cell type was defined by the maximum probability of cell type. This enabled us to visualize and dissect the complex spatial organization of various cell types present in the LUSC TME (**Figure 1c**). The number of cells analyzed for each sample is presented in **Supplementary** Figure 1a.

Integrated cell types were projected onto a UMAP plot ^27^, with each cell type identified by TACCO depicted in unique colors (**Figure 1c**). To validate these cell type annotations, UMAP plots were visualized with key markers such as EPCAM (epithelial), PECAM1 (endothelial), CD79A (B-cells), CD3D (T-cells), COL1A1 (fibroblasts), LYZ (macrophages), MARCO (macrophages), CD1C (dendritic cells), and NKG7 (NK cells) (illustrated in **Supplementary** Figure 1b). In addition, gene markers for each cell type were estimated from the MERFISH data by differentially expressed genes and are exhibited in **Figure 1d**. Markers derived from scRNA-seq data were also estimated and depicted, further confirming the cell types (**Supplementary** Figure 1c). Through this in-depth cell typing approach, we were able to demonstrate the extensive spatial heterogeneity of cell populations within the TME. Our spatial maps showed the variety of cell types and emphasized the potential linkages between their spatial distribution and the pathological characteristics of LUSC.

Cell type distribution across individual samples was evaluated and is presented in **Figure 2a**, revealing marked heterogeneity. Utilizing the spatial data from image-based ST, we analyzed the neighborhoods of these cell types^27^. Briefly, neighborhood analysis quantifies cell proximity within tissues, assigning high scores to closely neighboring cell types and low scores to distant ones. This was determined through a permutation test (**Figure 2b**). Whole slide images of H&E were assessed for the presence of immune infiltrates^28,29^. The visual immune subtypes were classified into three categories: immune-inflamed, immune-deficient, and intermediate. Notably, B cells, T cells, endothelial cells, and macrophages were found in significantly higher proportions in the ‘inflamed’ subtype, whereas epithelial cells were less prevalent (**Figure 2c**).

**Figure 2.**
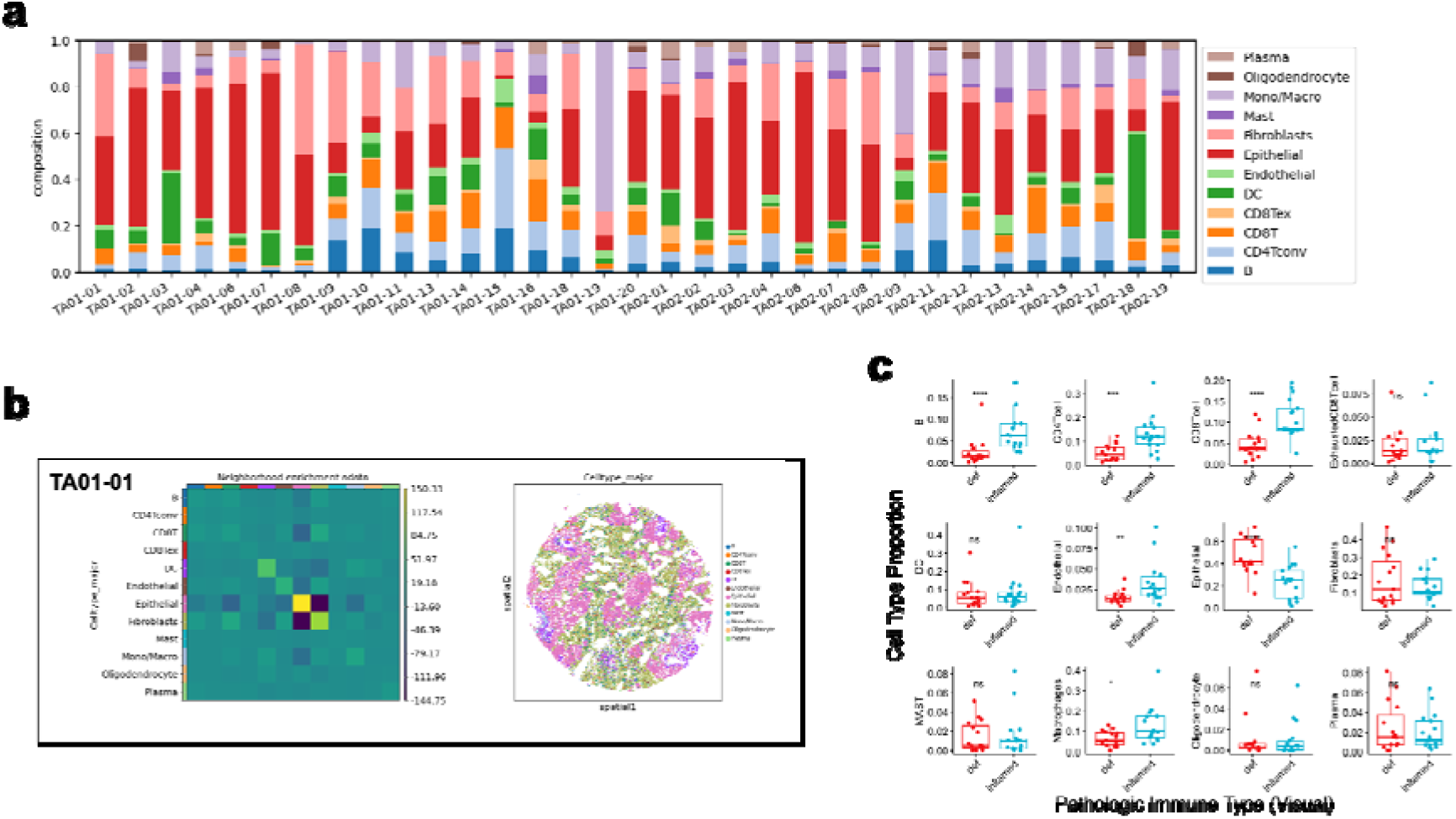
Analysis of cell type distribution and spatial relationship in LUSC TME. (a) Visualization of cell type heterogeneity across individual LUSC samples, highlighting the diverse distribution and abundance of cell types within the TME. (b) Results from a neighborhood analysis using spatial transcriptomics data, which quantifies the proximity of cell types within the tissue. High proximity scores indicate closely neighboring cell types, as determined by a permutation test, providing spatial relationship features reflecting cellular architecture of the TME. (c) Comparison of cell type proportions between pathological subtypes of LUSC, ‘inflamed’ and ‘immune deficient’. The analysis shows significant differences in the prevalence of B cells, T cells, endothelial cells, and macrophages between subtypes, illustrating the correlation between cellular composition and disease pathology.

### Tumor growth was associated with cancer associated fibroblasts and their spatial patterns in TME

We assessed the relationship between the proportions of cell types in the tumor microenvironment (TME) and key clinico-pathologic tumor characteristics — specifically, pathologic gross tumor size and maximum standardized uptake value (SUVmax) as measured by [^18^F]fluorodeoxyglucose PET imaging. These characteristics are crucial indicators of tumor growth and proliferative potential in lung cancer^30-32^. Spearman correlation values between these key characteristics and the proportions of cell types are depicted in **Figure 3a**. The results demonstrated a significant negative correlation between tumor size and the proportion of fibroblasts within the TME (rho = -0.44, p = 0.011), while the correlation between SUVmax and fibroblast proportion approached statistical significance (rho = -0.22, p = 0.22) (**Figure 3b**). This inverse relationship suggests that as tumors grow and become more proliferative, the fibroblast population within the TME decreases. These findings are corroborated by the analysis using data from both the Cancer Genome Atlas (TCGA) and the Cancer Imaging Atlas (TCIA)^33^. Tumor volume and TLRmax (SUVmax of tumor normalized by liver SUVmean) were analyzed with fibroblasts enrichment score estimated by bulk RNA-seq from TCGA^34^. This analysis revealed that TLRmax had a significant negative correlation with fibroblast enrichment scores (rho = -0.44, p = 0.046), while the correlation with tumor volume, although negative, did not reach statistical significance (rho = -0.33, p = 0.15) (**Supplementary** Figure 2).

**Figure 3.**
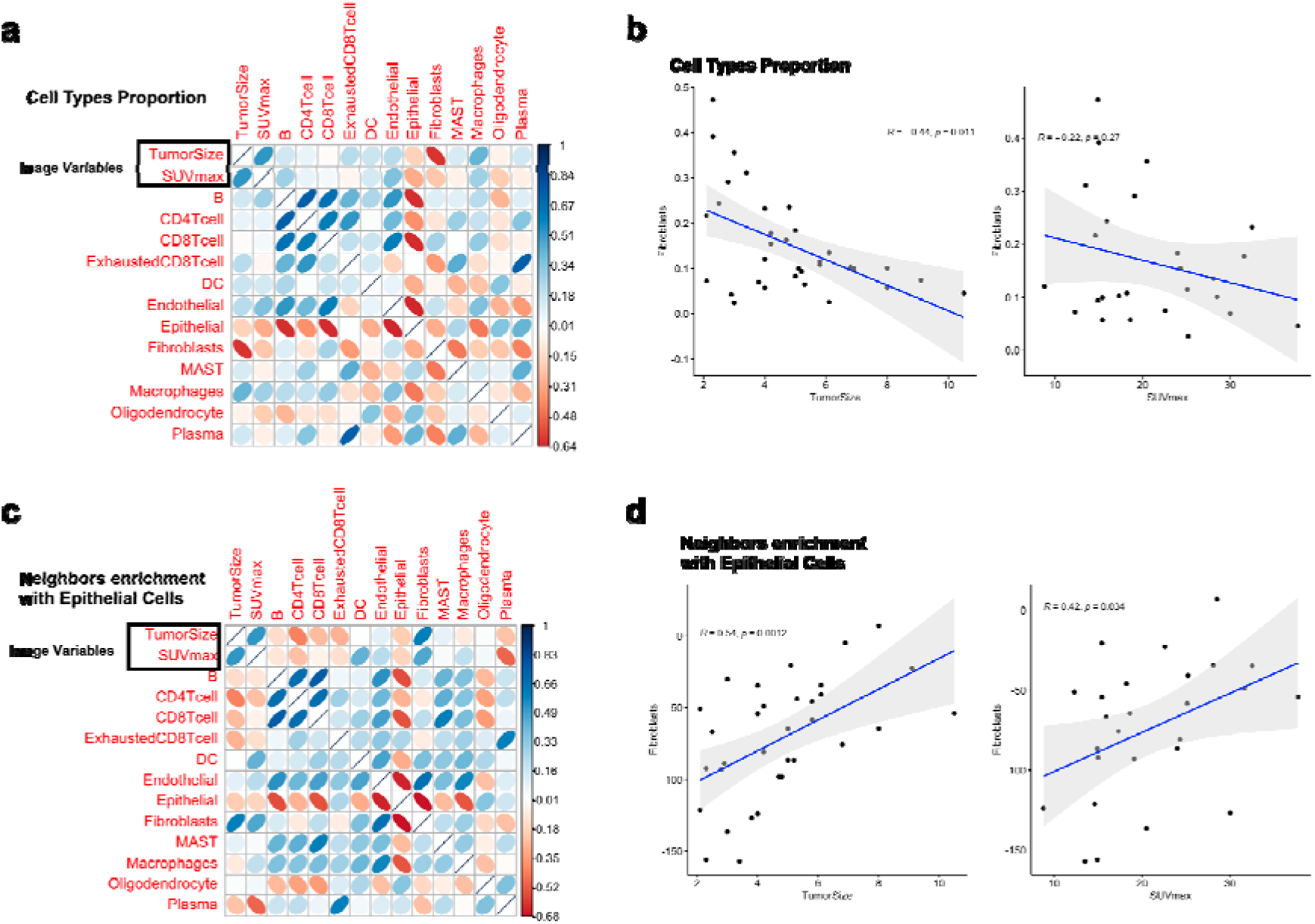
Correlations between cell type proportions or neighborhood enrichment in the TME and clinical tumor characteristics. (a) Spearman correlation analysis between key tumor features (pathologic gross tumor size and SUVmax) and the proportions of different cell types in the LUSC TME. This correlation plot presents correlation values, emphasizing relationships between tumor size and metabolic activity (SUVmax) and cellular composition in the TME. (b) Detailed correlation findings demonstrating a significant negative correlation between tumor size and fibroblast proportion, and a near-significant correlation with SUVmax measured on [^18^F]FDG PET (rho = -0.44, p = 0.01 for tumor size; rho = - 0.22, p = 0.27 for SUVmax). (c) Analysis of neighborhood enrichment scores calculated to assess the spatial proximity of tumor epithelial cells to other cell types, correlated with tumor size and SUVmax. This assessment helps understand the spatial dynamics within the TME in relation to tumor growth and metabolic activity. (d) Spearman correlation results for neighborhood enrichment scores between fibroblasts and epithelial cells in relation to tumor size and SUVmax. Notably, a significant positive correlation was found (rho = 0.54, p < 0.01 for tumor size; rho = 0.42, p = 0.03 for SUVmax).

Additionally, to explore the spatial dynamics of TME, we calculated the neighborhood enrichment scores between tumor epithelial cells and other cell types and examined their correlation with tumor size and SUVmax. These scores, particularly between ‘epithelial cells’ and other cell types, were used to assess spatial proximity to tumor epithelial cells within the TME. The spearman correlation analysis of these scores with tumor size and SUVmax is illustrated in **Figure 3c**. The neighborhood enrichment scores of fibroblasts and epithelial cells were presented with other clinical variables and the proportions of cell types in TME in **Supplementary** Figure 3. Notably, the neighborhood enrichment scores for fibroblasts exhibited a significant positive correlation with both tumor size (rho = 0.54, p = 0.001) and SUVmax (rho = 0.42, p = 0.03), as shown in **Figure 3d**. These results indicate that while the overall proportion of fibroblasts within the TME may decrease with tumor growth and gaining tumor proliferation features, the spatial closeness of fibroblasts to tumor epithelial cells increases.

### Proximity of fibroblasts to tumor epithelial cells associated with poor outcome in LUSC

The association of neighborhood enrichment scores of fibroblasts with tumor size and increased glucose metabolism in tumors led to further investigation into whether the proximity of fibroblasts to tumor epithelial cells influences LUSC prognosis. Utilizing the median neighborhood enrichment score as a threshold, LUSC patients were categorized into ‘Fibroblast-excluded’ and ‘Fibroblast-infiltrated’ groups. The ‘Fibroblast-infiltrated’ group exhibited poorer recurrence-free survival compared to the ‘Fibroblast-excluded’ group (**Figure 4a**). Notably, in **Figure 4b**, we analyzed the association between the proximity of tumor microenvironment cell types to tumor epithelial cells and recurrence free survivals. Hazard ratios (HRs) suggested that, aside from fibroblasts, which showed a significantly increased HR of 8.47 (95% CI: 1.06-67.79, p=0.044), and T staging with an HR of 9.09 (95% CI: 1.13-72.83, p=0.038). The proximity of other cell types to tumor epithelial cells were not associated with the prognosis in LUSC. As examples of cases with tumor microenvironment and gross tumor patterns, a ‘Fibroblast-infiltrated’ case demonstrated intense [^18^F]FDG uptake in the primary tumor (SUVmax = 22.6) with a tumor size of 8.7 cm, where spatial transcriptomics data revealed a mixed presence of tumor epithelium and fibroblasts. Conversely, a ‘Fibroblast-excluded’ example showed a clear separation of fibroblasts and tumor epithelial cells, with the patient having a smaller tumor (3.0 cm) and moderate [^18^F]FDG uptake (SUVmax = 13.5) (**Figure 4c**).

**Figure 4.**
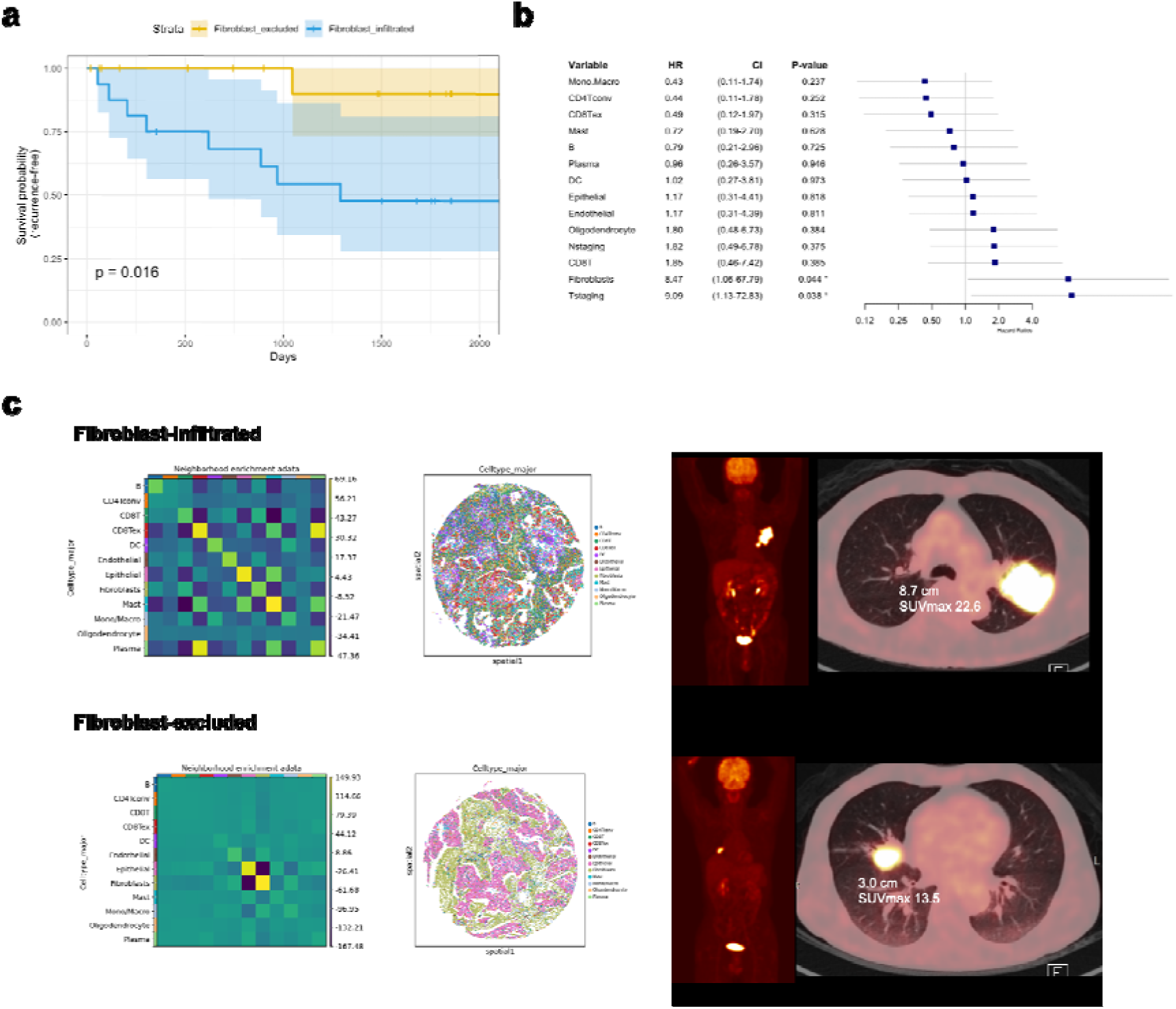
Prognostic impact of fibroblast proximity to tumor epithelial cells on LUSC. (a) Recurrence-free survival was analyzed LUSC patients categorized based on the median neighborhood enrichment score of fibroblasts. A Kaplan-Meier plot demonstrates that the ‘Fibroblast-infiltrated’ group has poorer survival outcomes compared to the ‘Fibroblast-excluded’ group (log-rank test, p = 0.02). (b) Analysis of the association between the proximity of various TME cell types to tumor epithelial cells and recurrence-free survival. Hazard ratios highlight the significant impact of fibroblast proximity on prognosis, with notable increases in risk, alongside T-staging influences on recurrence-free survival. (c) Representative cases showing the effect of fibroblast proximity on tumor metabolic activity and size. An example with ‘Fibroblast-infiltrated’ group shows high glucose metabolism and a larger tumor, whereas a ‘Fibroblast-excluded’ case exhibits lower metabolic activity and a smaller tumor size.

### Different states of fibroblasts according to spatial proximity to tumor epithelial cells

Considering the spatial dynamics between fibroblasts and tumor epithelium in LUSC, and its implications for prognosis, we further performed in-depth analysis of the molecular characteristics of fibroblasts relative to their spatial positioning. We delineated the tumor epithelial cell regions to measure the distance of fibroblasts from these areas using ST data (**Figure 5a**). The analysis revealed that the proximity to tumor epithelial regions influenced the distribution of various cell types within the TME. This analysis revealed patterns of TME cell types enrichment according to the positioning from tumor epithelial cells. Immune cells such as B-cells and CD8 T-cells densities increased with greater distances from the tumor epithelium, whereas exhausted CD8 T-cells predominated near the tumor epithelium and diminished with distance (**Figure 5b**). We categorized fibroblasts into two groups based on their median distance from tumor epithelium: Epithelial-distant fibroblasts and epithelial-adjacent fibroblasts, and analyzed their molecular markers (**Figure 5c**). Markers such as CXCL12, MMP2, MZB1, and PDGFRA were characteristic of epithelial-distant fibroblasts and were prevalent in the central zones of fibroblast-rich TME regions in LUSC samples (**Figure 5c, Supplementary** Figure 4). Conversely, epithelial-adjacent fibroblasts exhibited markers like MMP11, PDGFRB, CCL26, and CD248, predominantly expressed at the peripheries of fibroblast-rich regions (**Figure 5c, Supplementary** Figure 4).

**Figure 5.**
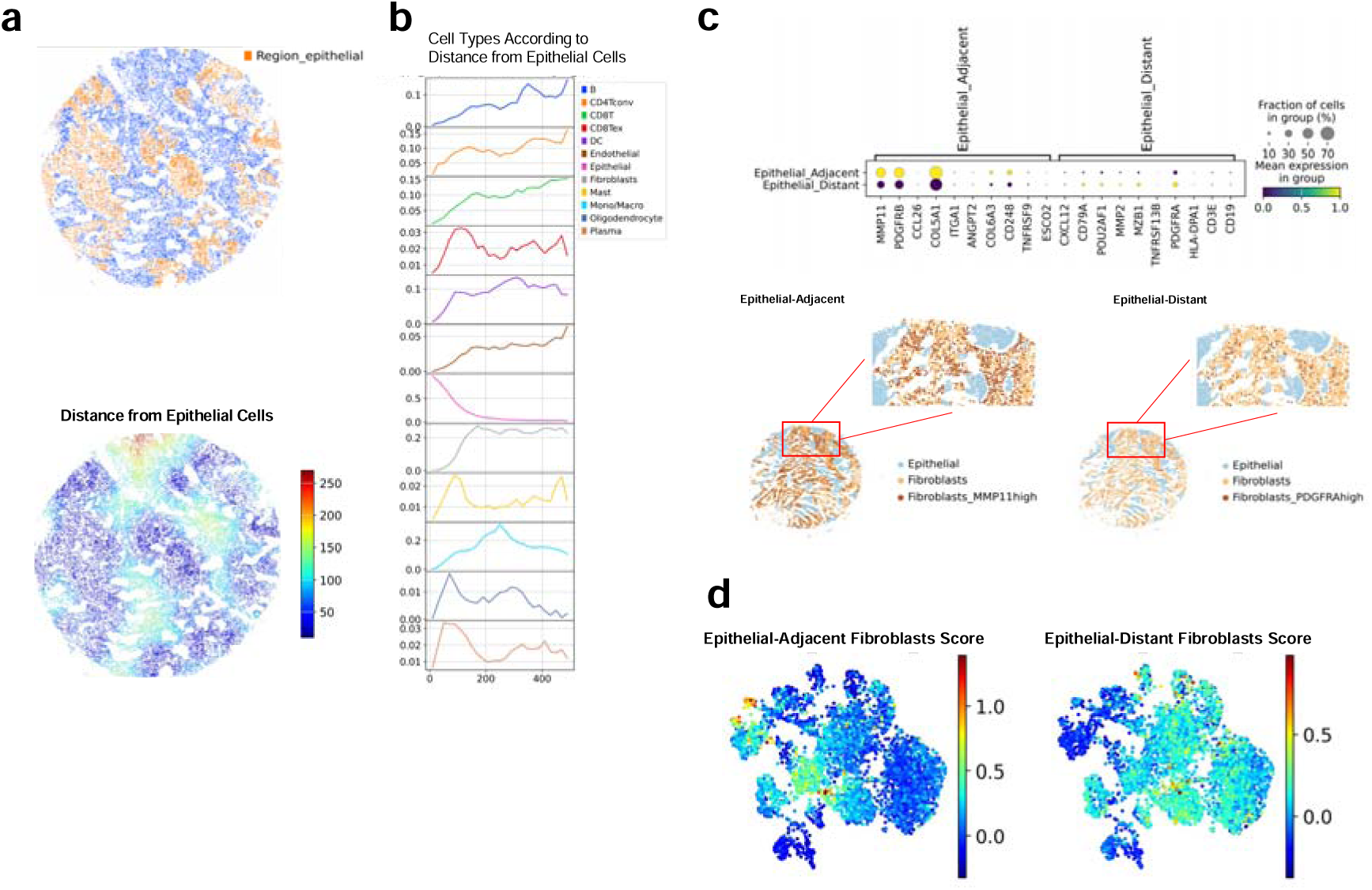
Molecular characterization of fibroblasts in relation to their spatial positioning within the TME. (a) Visualization of fibroblast distribution relative to tumor epithelial regions using spatial transcriptomics data. The spatial distances of cells from epithelial areas were measured to analyze the molecular features of spatial positioning on their distribution within the TME. (b) Exploration of cell type enrichment patterns in relation to their distance from tumor epithelial cells. The figure shows increased densities of immune cells like B-cells and CD8 T-cells at greater distances from the tumor epithelium, whereas exhausted CD8 T-cells are more prevalent closer to the epithelium and decrease with distance. (c) Categorization and molecular characterization of fibroblasts into ‘Epithelial-adjacentt’ and ‘Epithelial-distant’ groups based on their median distances from tumor epithelium. Specific molecular markers, including MMP11 for epithelial-adjacent fibroblasts and PDGFRA for epithelial-distant fibroblasts, are highlighted to show their distinct molecular profiles depending on their spatial positioning. (d) Enrichment scores for molecular signatures of epithelial-distant and epithelial-adjacent fibroblasts analyzed on cancer-associated fibroblasts (CAFs) subset of scRNA-seq data. The spatial proximity of fibroblasts to tumor epithelial cells correlates with distinct molecular signatures, which in turn are associated with specific subtypes of CAFs.

To investigate deeper into the markers of fibroblasts, we further analyzed fibroblasts from scRNA-seq data (GSE131907). We isolated a subset of fibroblasts and clustered them based on their transcriptomic profiles (**Supplementary** Figure 5a). The key markers of epithelial-distant and epithelial-adjacent fibroblasts were represented on the UMAP of scRNA-seq data (**Supplementary** Figure 5b). Moreover, we calculated and visualized the enrichment scores for the signatures of epithelial-distant and epithelial-adjacent fibroblasts within the scRNA-seq dataset (**Figure 5d, Supplementary** Figure 5c), which also delineated distinct clusters. Drawing on prior pan-cancer research on CAF states^6^, we visualized key characteristic features of three major CAF states, myofibroblasts, inflammatory fibroblasts, and adipogenic signals (**Supplementary** Figure 5d). Additionally, ‘cluster 4’ and ‘cluster 5’ exhibited high scores for the epithelial-adjacent fibroblasts signature, whereas ‘cluster 0’ and ‘cluster 1’ demonstrated high scores for the epithelial-distant fibroblasts signature (**Supplementary** Figure 5e). The primary markers of these clusters are detailed in **Supplementary Table 2**. Specifically, IGFBP6+ CAFs were categorized within ‘cluster 0’, and POSTN+ myofibroblasts were categorized within ‘cluster 4’. Our analysis indicated an association between the epithelial-adjacent fibroblast signature and the myo- and inflammatory CAF states, whereas the epithelial-distant fibroblast signature was linked to adipogenic CAF states.

To elucidate the crucial molecular characteristics linked to the interaction between fibroblasts and tumor epithelial cells, we extended our analysis to the whole transcriptome level. Given the limitations of image-based ST in terms of gene panel size, we incorporated barcode-based ST (Visium, 10x genomics) to comprehensively assess gene expression in LUSC while retaining spatial context. We acquired Visium data from tumor samples of eight LUSC patients. **Supplementary Table 3** summarizes their brief demographics and clinical details. Using CellDART^35^ with the reference scRNA-seq data, we mapped the spatial distribution of cell types within the TME, with a focus on epithelial cells and fibroblasts (**Figure 6a**; complete cell type mapping results are available in **Supplementary** Figure 6). To uncover molecular patterns in regions where fibroblasts and epithelial cells spatially coincide, we analyzed topologically overlapping cell type patterns in the barcode-based ST data^36^. This analysis highlighted regions with a high density of specific cell types and their intersections with other cell populations (**Figure 6b**). Specifically, we compared regions where fibroblasts were enriched and overlapped with epithelial cells to those where fibroblasts were distant from epithelial cells (**Figure 6c**). We identified key markers and conducted gene ontology analysis for these regions (**Figure 6d**). Intriguingly, areas where fibroblasts overlapped with epithelial cells showed a strong association with the glycolysis process (**Figure 6d**), aligning with the active glucose uptake observed in [^18^F]FDG PET imaging in LUSC with high neighborhood enrichment of fibroblasts.

**Figure 6.**
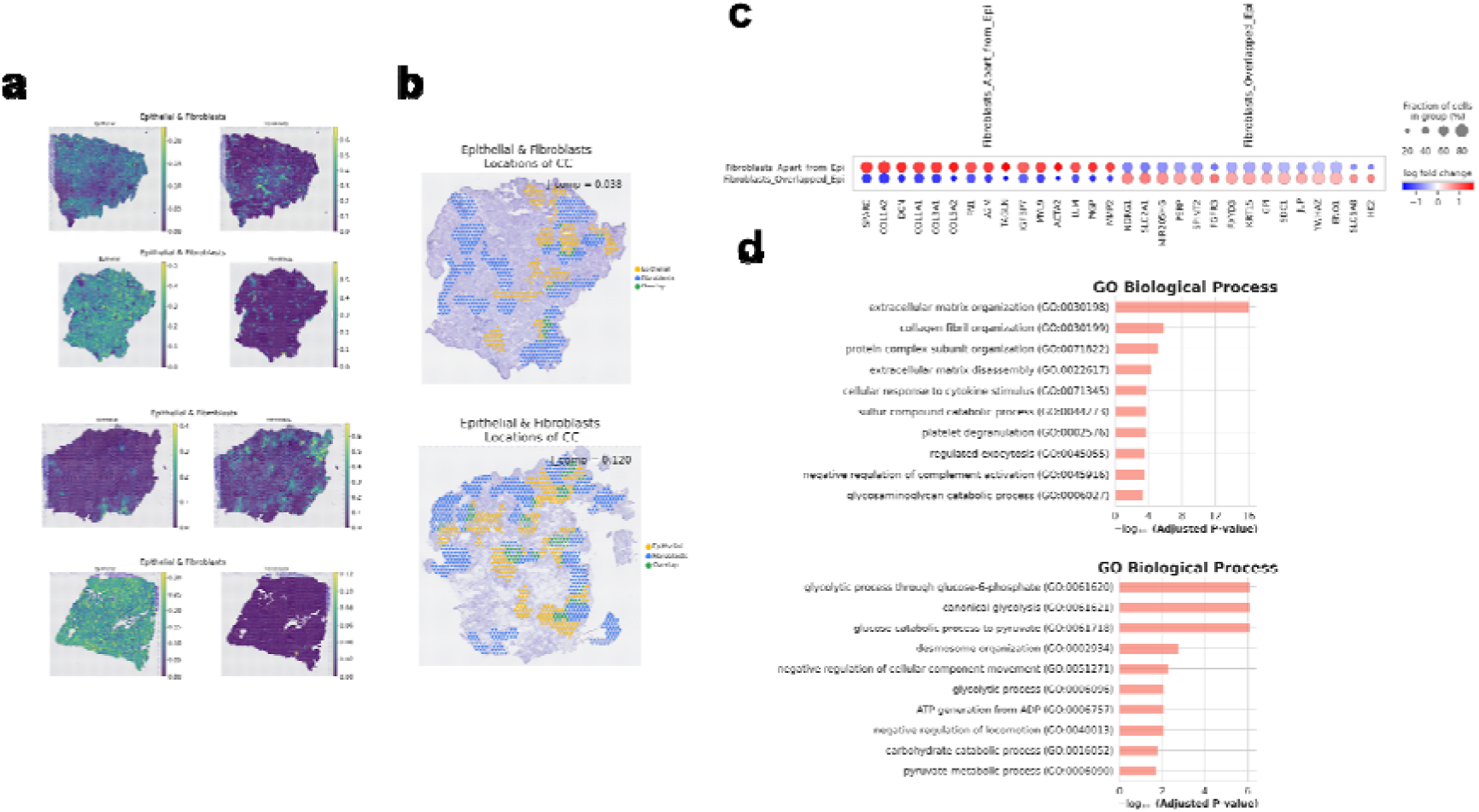
Topological overlap patterns of fibroblast-epithelial cells in the TME. (a) Visualization of the spatial distribution of cell types within the LUSC TME using barcode-based spatial transcriptomics (Visium). Two sample images highlights the detailed cell type mapping of epithelial cells and fibroblasts. (b) Topologically overlapping cell type patterns were analyzed in the barcode-based spatial transcriptomics data. Regions with a locally high density of fibroblasts are indicated in blue, while areas with a high density of epithelial cells are shown in yellow. Regions where these cell types overlap are marked in green. (c) Identification of key markers for regions of fibroblast and epithelial cell overlap versus fibroblast-rich regions without overlapping. (d) Gene ontology analysis for key markers of regions of fibroblast and epithelial cell overlap was performed. Notably, areas of overlap are associated with enhanced glycolysis processes, reflecting the active metabolic interactions that correspond to the observed glucose uptake patterns in LUSC as detected by [^18^F]FDG uptake according to the proximity of fibroblasts to tumor epithelium.

## Discussion

Our research advances the understanding of the spatial complexity of CAFs in LUSC, highlighting their profound influence on tumor growth and patient outcomes. CAFs have been increasingly recognized for their critical role in structuring the TME and influencing the response to treatments, especially in immuno-oncology^8-12^. Given their heterogeneity and diverse roles in promoting tumor growth and providing support through various protumoral factors, CAFs have emerged as potential therapeutic targets in the stromal components of the TME across various solid tumors^3,37,38^. Despite the spatial diversity of CAFs, ranging from their infiltration into the tumor core to their prevalence in stroma-rich peripheral regions, a comprehensive understanding of their spatial patterns, molecular characteristics, and their association with clinical parameters like tumor growth and prognosis remains elusive^39-41^. While scRNA-seq has shed light on the molecular features and subtypes of CAFs, as well as their complex interactions with other cells, particularly tumor cells, the spatial dynamics of these heterogeneous CAF populations have been less explored due to the challenges in capturing their spatial context particularly in lung cancer. Through the application of ST, our study investigated the spatial organization of CAFs within the TME and its association with clinical outcome as well as tumor size and glucose metabolism. By integrating spatial and molecular data, we revealed how the localization and molecular signatures of CAFs interact with tumor dynamics, providing valuable insights that could inform targeting the spatial heterogeneity of CAFs and their architecture of TME.

Our study elucidates the role of fibroblast proximity to tumor epithelial cells in tumor growing and obtaining aggressiveness reflected by clinical outcomes LUSC. By distinguishing fibroblasts as ‘Fibroblast-infiltrated’ or ‘Fibroblast-excluded’ based on their neighborhood enrichment score, we discovered that patients with greater fibroblast infiltration near tumor epithelial cells tend to have poorer recurrence-free survival. This observation aligns with our findings that the spatial enrichment of fibroblasts relative to tumor epithelial cells correlates significantly with tumor size and glucose metabolism clinically estimated by [^18^F]FDG PET imaging. This correlation suggests a dynamic reorganization of CAFs in response to tumor progression—from a state of spatial separation from the tumor epithelium to one of increased infiltration^42^. This dynamic spatial distribution of CAFs is crucial for understanding tumor behavior and patient prognoses, reinforcing the significance of the spatial architecture within the TME. The significance of this spatial reorganization aligns with findings from a prior study that demonstrated the prognostic relevance of pathologic patterns in LUSC, specifically highlighting cases where carcinoma cells are surrounded by fibrous stroma as independent predictors of patient outcomes^8,43^. Our findings thus highlight the critical nature of CAF spatial organization in influencing tumor progression and underscore the potential of biomarkers and targeting spatial dynamics to improve LUSC patient outcomes.

In addition to linking fibroblast spatial patterns with clinical outcomes, we unveiled unique molecular signatures for fibroblast subpopulations based on their proximity to tumor epithelial cells. Epithelial-distant fibroblasts, identified by markers like CXCL12 and PDGFRA, predominated in the core regions of fibroblast-dense TME zones. In contrast, epithelial-adjacent fibroblasts, distinguished by markers such as MMP11 and PDGFRB, were mainly found at the edges of these zones. This delineation underscores that a spatial position of fibroblasts relative to tumor epithelial cells might dictate its functional role within the TME. Notably, the differential expression of PDGFRA and PDGFRB, crucial CAF markers, according to fibroblast subtypes in immunohistochemistry analyses, aligns with a previous report where PDGFRB-positive CAFs were linked with tumor epithelial-adjacent fibroblasts^44^. Furthermore, our analysis identified an ‘epithelial-adjacent’ signature correlating with inflammatory and myofibroblastic CAF subtypes previously characterized in pan-cancer studies, while the ‘epithelial-distant’ signature aligned with adipogenic CAF traits^6^. Expanding on the significance of inflammatory CAFs (iCAFs), these cells have been implicated in various critical tumor-promoting processes, from remodeling the extracellular matrix to modulating the immune landscape within the TME^2^. They are known to secrete a range of cytokines, chemokines, and growth factors that can recruit immune cells and support tumor growth and progression. Our findings related to this paradigm, suggesting that the proximity of CAFs to tumor cells might enhance their ability to influence the tumor immune microenvironment and potentially its response to immunotherapy. Notably, PDGFR and MMP, which belong to the same class of molecules associated with CAFs or CAF-secreted matrix metalloproteinases, exhibit varying types based on their proximity to the tumor epithelium. According to our results, PDGFRB^+^MMP11^+^ CAFs were found close to the tumor epithelium. This spatial position could affect immune profiles of tumor epithelium as secreting factors such as MMP11, which is acting as a potential immune modulator that may contribute to resistance to immunotherapy in previous studies^45,46^. Accordingly, exploring the spatial dynamics of CAFs further, particularly subtypes of CAFs, could pave the way for the development of novel spatial biomarkers supporting the intricate roles of CAFs in modulating the immune landscape of tumors.

The observed association between increased tumor glycolysis activity and the proximity of fibroblasts to tumor epithelium underscores the influential role of CAFs in shaping tumor metabolism. Our study investigates the molecular characteristics of the regions where fibroblasts and epithelial cells intersect in LUSC, discovering a significant association with glycolytic processes. This correlation is also supported by active glucose uptake, as evidenced by [^18^F]FDG PET imaging associated with neighborhood of fibroblasts to tumor epithelium increased according to [^18^F]FDG uptake. Previous studies have showed that CAFs can modulate cancer cell metabolism through multiple mechanisms^47^. CAFs supply metabolic byproducts that allow cancer cells to allocate glucose towards anabolic pathways, essential for their rapid proliferation^48,49^. Moreover, CAFs can enhance the expression of glucose transporters on tumor cells through a variety of secreted factors and direct interactions, thereby boosting glucose absorption^50^. By reshaping metabolic profiles of the TME, CAFs set the stage for tumor advancement and aggression. Their activity contributes to a hypoxic environment that further alters tumor cell metabolism and fosters more malignant behaviors^51,52^. Our ST-based analysis, revealing the interplay between CAFs and tumor epithelium in LUSC, sheds light on the glycolytic enhancements within these interactive regions. This insight into the metabolic crosstalk between fibroblasts and cancer cells opens up new avenues for identifying metabolic vulnerabilities that could be targeted therapeutically in LUSC.

Our study leveraged the strengths of both image-based (MERFISH) and barcode-based ST (Visium) platforms to examine the spatial relationships of cell types within the TME of LUSC. These platforms allowed us to transcend the limitations of gene panel size of image-based ST and spatial resolution inherent to barcode-based ST methodologies. By providing a detailed spatial and molecular landscape of LUSC, our research offers a more comprehensive understanding of the TME, which is crucial for identifying new biomarkers and therapeutic targets by particularly characterizing CAFs according to the spatial context in the TME. However, a limitation of our study is the separate utilization of ST data from different samples, which precluded a direct comparison of the datasets. Future research, focusing on simultaneous spatial and molecular analyses using ST data from the same tumor samples, will likely yield deeper insights into the spatial dynamics and functional roles of CAFs within the actual human tumor setting.

In conclusion, our research underscores the critical role of spatial organization and molecular features of CAFs in influencing LUSC progression. The insights gained from this study highlight the potential of ST in unraveling the complex spatial dynamics of the TME, offering new avenues for targeted therapies that consider the spatial context of cellular interactions within tumors.

## Methods

### Tissue preparation and MERSCOPE

A tissue microarray (TMA) was constructed from surgically resected LUSC specimens preserved as formalin-fixed, paraffin-embedded (FFPE) tissues, featuring 2 mm cores. Selection of specific core locations within the whole FFPE blocks was guided by a thorough examination of H&E stained whole slide images from each tumor section reviewed by pathologists. The creation of the TMA was in compliance with all protocols approved by the Institutional Review Board (IRB Application No.: H-2009-081-1158).

For the MERFISH analysis using the MERSCOPE system (Vizgen Inc., MA, USA), we chose a 500-gene panel, which includes pre-selected markers crucial for identifying various cell types within the TME. Details of the gene panel can be found in Supplementary Table 3. RNA extraction was conducted using the PureLink™ FFPE RNA Isolation Kit (Invitrogen, Waltham, MA), adhering to the manufacturer’s protocol. The quality of the RNA was evaluated by DV200, the percentage of RNA fragments larger than 200 nucleotides relative to all RNA fragments, using the Agilent RNA 6000 Pico Kit The TMA tissue sections were placed on glass coverslips designed for the MERSCOPE system. Tissue processing and imaging adhered to the MERSCOPE-recommended protocols for FFPE samples. The procedure initiated with deparaffinization the tissue on the MERSCOPE slide for dissolving the paraffin, followed by decrosslinking for 15 minutes at 95°C. The tissue was incubated in the Pre-Anchoring Reaction Buffer for 2 hours at 37°C and then overnight in the Anchoring Buffer at 37°C. The subsequent day involved gel embedding and clearing ranging from 68 hours, aligning with the guidelines for resistant FFPE tissues outlined by MERSCOPE. After clearing, the tissue was hybridized with the encoding probe for 45 hours, followed by staining with DAPI and PolyT Staining Reagent for 15 minutes. Finally, the MERSCOPE instrument setup was completed, and tissue imaging was conducted as per the instructions in the Vizgen MERSCOPE Instrument User Guide (https://vizgen.com/resources/user-guides/).

### Tissue preparation and barcode-based ST using Visium

LUSC specimens were obtained from eight patients who underwent surgical resection. In the operating room, these specimens were immediately embedded in OCT compound and subsequently stored at -80°C until needed for cryosectioning for Visium (10X genomics, USA). For this process, samples were brought to -20°C using a cryotome (Thermo Scientific, USA), and sections were prepared for spatially resolved gene expression analysis employing the Visium Spatial Transcriptomic kit (10X Genomics, USA). The procedure began with incubating the tissue sections with permeabilization enzymes, the duration of which was optimized using Visium Spatial Optimization Slides (1000193, 10X Genomics). After a saline sodium citrate buffer wash, the sections were treated with RT Master Mix (from the Visium Reagent kit, 10X Genomics) to carry out reverse transcription as per the kit’s instructions. Following reverse transcription, the sections were exposed to 0.08 M KOH for five minutes and then to Second Strand Mix for fifteen minutes at 65°C, preparing them for cDNA amplification and library construction. Sequencing of the libraries was executed using the NovaSeq 6000 System S1 200 (Illumina, CA, USA) aiming for a sequencing depth of around 250 million read pairs per sample. For gene expression analysis, raw FASTQ data along with H&E-stained images were processed using Space Ranger v1.1.0 software (10X Genomics). The ‘mkfastq’ command facilitated the conversion of Illumina base call files to FASTQ format for individual samples. The ‘count’ command was employed to analyze Visium spatial expression libraries, which involved aligning the images to the fiducial alignment grid to ascertain the orientation and position of the tissue sections. The STAR (v2.5.1b) aligner was utilized to align sequencing reads, and gene expression in each tissue section spot was quantified using the unique molecular identifier and 10X barcode data.

### Processing of scRNA-seq data

The publicly available single-cell RNA sequencing data of NSCLC, from dataset GSE131907, were utilized to map cell types^7^. These cell types were mapped using a pre-processed version of a uniform analysis pipeline curated by the Tumor Immune Single-cell Hub2 (TISCH2), which was downloaded from an available source^53^. Visualization of these cell types was achieved using UMAP as a dimensionality reduction method. Cell type markers were identified using the scanpy module (version 1.9.6)^54^, employing the Wilcoxon rank-sum method.

To explore specific subsets of fibroblasts within this dataset, fibroblast cells were isolated and subjected to further clustering using the Leiden clustering method (scanpy.tl.leiden). Accordingly, 4125 cells labeled with fibroblasts were further analyzed. By adjusting the resolution parameter (resolution = 0.3), 10 distinct clusters of cell types were identified, and their markers were determined. For visualizing the enrichment scores of ‘epithelial-adjacent fibroblasts’ and ‘epithelial-distant fibroblasts,’ as estimated by spatial transcriptomics, module scores were computed using the ‘scanpy.tl.score_genes’ function. This function calculates the average expression of a set of genes minus the average expression of a reference gene set.

### Processing and Cell typing of MERFISH data

Image-based ST data, obtained using MERSCOPE, were partitioned into individual samples from a tissue microarray. For computational analysis, TMAtools software (Portrai Inc., Republic of Korea) was employed, which automatically segregated the core of TMA-based ST data. Following the segregation, cell types were identified by integrating with scRNA-seq data, which had previously been labeled with cell types (as described in the previous section). TACCO algorithm was utilized for cell type labeling, using its default settings^26^. The ‘probability’ of cell types for each cell in image-based ST was stored in ‘AnnData.obsm’. For deterministic cell type identification, the cell type with the highest probability was selected as the definitive cell type for each cell.

To integrate the different cores of image-based ST data, batch correction was performed using the scVI algorithm^55^. Considering potential batch effects from various samples, ‘different cores’ of the TMA were utilized as batch covariates in the integration process using scVI tools. The parameters set for this integration included: number of latent dimensions = 50, number of layers = 3, dropout rate = 0.1, and batch size = 1024. UMAP was then used to visualize the two-dimensional gene expression data space of cells in the image-based ST data. After cell type labeling with TACCO, cell type markers for the image-based ST data were also identified using Wilcoxon rank-sum tests.

### Neighborhood enrichment score estimation on image-based ST data

Neighborhood enrichment analysis was conducted using cell types and their spatial locations from image-based ST data. This analysis involved calculating an enrichment score that measured the proximity of cell types within a connectivity graph. The observed number of events was then compared to a series of permutations, and a z-score was computed to assess statistical significance using Squidpy (specifically, the function squidpy.gr.nhood_enrichment) (version 1.3.1)^56^.

### Analysis for spatial relationship of TME cell types

Spatial distance-based analysis was conducted using the epithelial regions of each core. Tumor epithelial regions were simply defined by identifying epithelial cell types, and distances from these cells were measured. The density of each cell type was then quantified based on its distance from the tumor epithelial regions. This process was based on TACCO package.

To further characterize fibroblasts based on their proximity to tumor epithelial regions, fibroblasts from image-based ST data were categorized into two subtypes using the median distance from tumor epithelial regions: ‘Epithelial-adjacent fibroblasts’ and ‘Epithelial-distant fibroblasts’. Additionally, an analysis of differentially expressed genes was performed to understand the characteristics of these two fibroblast subtypes. The top 100 genes enriched in fibroblasts from the panel of 500 genes were selected based on fold-change of fibroblasts. A Wilcoxon rank sum test was used to evaluate differential expression among these top 100 genes to find gene features of two fibroblast subtypes in relation to their spatial proximity to tumor epithelial cells.

### Cox proportional hazard model for recurrence free survival

Among 33 patients with LUSC, we divided the patients into two groups based on the proximity of specific cell types to tumor epithelial cells. These binary subgroups were then analyzed using a Cox proportional hazard model to estimate the hazard ratio and determine the statistical significance of variables associated with recurrence-free survival. This analysis was limited to univariate analysis. Additionally, pathologic T-staging and N-staging were evaluated for comparison. For the proximity of fibroblasts to tumor epithelial cells, patients were categorized into two subtypes: ‘Fibroblast-infiltrated’ and ‘Fibroblast-excluded’, based on the median value of neighborhood enrichment scores. Kaplan-Meier curves for these subgroups were plotted to assess recurrence-free survival, and a log-rank sum test was performed.

### Pathological immune type of LUSC

The pathological immune subtypes of lung squamous cell carcinoma (LUSC) samples were assessed by lung cancer pathology experts through visual analysis. H&E stained sections of resected lung specimen of cancer patients were assessed for the presence of immune infiltrates as previous paper with slight modification^28,29^. Briefly, the extent of immune cell infiltration in whole section of tumor tissue was assessed. The cases with no to minimal immune cells infiltration were classified as “immune deficient”, and cases with moderate to marked immune cells infiltration were classified as “immune inflamed”.

### [^18^F]FDG PET data and imaging features of LUSC patients

According to the standard protocol of the hospital, patients were required to fast for at least six hours before being administered 5.18 MBq/kg of [^18^F]FDG intravenously. PET/CT scans were acquired 60 minutes after the injection (Biograph 40 and Biograph 64, mCT, Siemens). An emission scan covering the area from the skull base to the proximal thigh was performed, accompanied by a CT scan for attenuation correction. All PET images were analyzed by an experienced nuclear medicine physician using commercial imaging software (Syngo.via, Siemens Healthcare, Erlangen, Germany). Spherical volumes of interest (VOIs) were delineated for each patient to determine the maximum standardized uptake value (SUVmax).

For the analysis combining TCGA data with TCIA data, TCGA-LUSC samples were selected, specifically those with corresponding [^18^F]FDG PET data from TCIA. The [^18^F]FDG uptake in tumors was assessed using the tumor-to-liver uptake ratio (TLRmax) due to the heterogeneous nature of data acquisition across different centers and protocols in the TCIA. The methods for this analysis are detailed in a previous study^57^. The tumor microenvironment cell types were evaluated using xCell analysis^34^, with a specific focus on the correlation between fibroblast enrichment scores, TLRmax, and tumor volume using Spearman’s correlation.

### Cell typing for barcode-based ST data and topological overlap analysis of cell types

In barcode-based ST data, cell types for each spot were deconvoluted by CellDART algorithm, referencing the previously mentioned scRNA-seq data. This analysis involved configuring CellDART with specific parameters: 80,000 pseudospots, 3,000 iterations, and 10 mixed cells per spot^35^. Subsequently, we utilized all pairs of cell type scores from these spots for topological colocalization analysis via STopover, applying default settings to assess the topological overlap patterns for each cell type within the barcode-based ST data^36^.

To explore the spatial co-localization patterns of fibroblasts with epithelial cells, we analyzed differentially expressed genes between regions defined by STopover. Notably, barcode-based ST data encompasses all genes, unlike image-based ST data. Additionally, the overlapping regions between fibroblasts and epithelial cells identified through this analysis do not necessarily define the cell types themselves, as they are derived from deconvolution results of the spots. Thus, the analysis of differentially expressed genes helps characterize spatial regions where epithelial cells and fibroblasts are highly overlapped. For differentially expressed genes, Wilcoxon rank sum tests were used. Gene ontology analysis was additionally conducted using gene ontology biological process terms using gseapy^58^.

## Acknowledgements

*Author’s contributions:* H.C. and Y.T.K. conceived the study. K.J.N. and Y.T.K. designed the clinical study and collected clinical samples and data. Y.J., M.L., J.E.L., and H.J.I. contributed to generate the data and perform experiments. H.C., K.J.N., Dj.L., and J.K. analyzed the data. H.C., K.J.N., J.K., and Y.T.K. interpreted clinical results. All authors contributed to the interpretation of the data and wrote the paper.

## References

1 Mascaux, C. et al. Immune evasion before tumour invasion in early lung squamous carcinogenesis. Nature 571, 570–575 (2019).

2 Sahai, E. et al. A framework for advancing our understanding of cancer-associated fibroblasts. Nature Reviews Cancer 20, 174–186 (2020).

3 Chen, Y., McAndrews, K. M. & Kalluri, R. Clinical and therapeutic relevance of cancer-associated fibroblasts. Nature reviews Clinical oncology 18, 792–804 (2021).

4 Lavie, D., Ben-Shmuel, A., Erez, N. & Scherz-Shouval, R. Cancer-associated fibroblasts in the single-cell era. Nature cancer 3, 793–807 (2022).

5 Hanley, C. J. et al. Single-cell analysis reveals prognostic fibroblast subpopulations linked to molecular and immunological subtypes of lung cancer. Nature communications 14, 387 (2023).

6 Luo, H. et al. Pan-cancer single-cell analysis reveals the heterogeneity and plasticity of cancer-associated fibroblasts in the tumor microenvironment. Nature communications 13, 6619 (2022).

7 Kim, N. et al. Single-cell RNA sequencing demonstrates the molecular and cellular reprogramming of metastatic lung adenocarcinoma. Nature communications 11, 2285 (2020).

8 Bremnes, R. M. et al. The role of tumor stroma in cancer progression and prognosis: emphasis on carcinoma-associated fibroblasts and non-small cell lung cancer. Journal of thoracic oncology 6, 209–217 (2011).

9 Wald, O. et al. Interaction between neoplastic cells and cancer-associated fibroblasts through the CXCL12/CXCR4 axis: Role in non–small cell lung cancer tumor proliferation. The Journal of thoracic and cardiovascular surgery 141, 1503–1512 (2011).

10 Navab, R. et al. Prognostic gene-expression signature of carcinoma-associated fibroblasts in non-small cell lung cancer. Proceedings of the National Academy of Sciences 108, 7160–7165 (2011).

11 Abulaiti, A. et al. Interaction between non-small-cell lung cancer cells and fibroblasts via enhancement of TGF-β signaling by IL-6. Lung cancer 82, 204–213 (2013).

12 Xiang, H. et al. Cancer-associated fibroblasts promote immunosuppression by inducing ROS-generating monocytic MDSCs in lung squamous cell carcinoma. Cancer immunology research 8, 436–450 (2020).

13 Mao, X. et al. Crosstalk between cancer-associated fibroblasts and immune cells in the tumor microenvironment: new findings and future perspectives. Molecular cancer 20, 1–30 (2021).

14 Jenkins, L. et al. Cancer-associated fibroblasts suppress CD8+ T-cell infiltration and confer resistance to immune-checkpoint blockade. Cancer research 82, 2904–2917 (2022).

15 Lakins, M. A., Ghorani, E., Munir, H., Martins, C. P. & Shields, J. D. Cancer-associated fibroblasts induce antigen-specific deletion of CD8+ T Cells to protect tumour cells. Nature communications 9, 948 (2018).

16 Rao, A., Barkley, D., França, G. S. & Yanai, I. Exploring tissue architecture using spatial transcriptomics. Nature 596, 211–220 (2021).

17 Ståhl, P. L. et al. Visualization and analysis of gene expression in tissue sections by spatial transcriptomics. Science 353, 78–82 (2016).

18 Williams, C. G., Lee, H. J., Asatsuma, T., Vento-Tormo, R. & Haque, A. An introduction to spatial transcriptomics for biomedical research. Genome Medicine 14, 68 (2022).

19 Rodriques, S. G. et al. Slide-seq: A scalable technology for measuring genome-wide expression at high spatial resolution. Science 363, 1463–1467 (2019).

20 Vickovic, S. et al. High-definition spatial transcriptomics for in situ tissue profiling. Nature methods 16, 987–990 (2019).

21 Chen, A. et al. Spatiotemporal transcriptomic atlas of mouse organogenesis using DNA nanoball-patterned arrays. Cell 185, 1777–1792. e1721 (2022).

22 Chen, K. H., Boettiger, A. N., Moffitt, J. R., Wang, S. & Zhuang, X. Spatially resolved, highly multiplexed RNA profiling in single cells. Science 348, aaa6090 (2015).

23 Janesick, A. et al. High resolution mapping of the tumor microenvironment using integrated single-cell, spatial and in situ analysis. Nature Communications 14, 8353 (2023).

24 He, S. et al. High-plex imaging of RNA and proteins at subcellular resolution in fixed tissue by spatial molecular imaging. Nature Biotechnology 40, 1794–1806 (2022).

25 Anderson, A. C. et al. Spatial transcriptomics. Cancer Cell 40, 895–900 (2022).

26 Mages, S. et al. TACCO unifies annotation transfer and decomposition of cell identities for single-cell and spatial omics. Nature biotechnology 41, 1465–1473 (2023).

27 Becht, E. et al. Dimensionality reduction for visualizing single-cell data using UMAP. Nature biotechnology 37, 38–44 (2019).

28 Montironi, C. et al. Inflamed and non-inflamed classes of HCC: a revised immunogenomic classification. Gut 72, 129–140 (2023).

29 Sia, D. et al. Identification of an immune-specific class of hepatocellular carcinoma, based on molecular features. Gastroenterology 153, 812–826 (2017).

30 Vesselle, H. et al. Lung cancer proliferation correlates with [F-18] fluorodeoxyglucose uptake by positron emission tomography. Clinical Cancer Research 6, 3837–3844 (2000).

31 Higashi, K. et al. FDG PET measurement of the proliferative potential of non-small cell lung cancer. Journal of Nuclear Medicine 41, 85–92 (2000).

32 Vesselle, H. et al. Relationship between non-small cell lung cancer FDG uptake at PET, tumor histology, and Ki-67 proliferation index. Journal of Thoracic Oncology 3, 971–978 (2008).

33 Kirk, S., Lee, Y. & Kumar, P. The cancer genome atlas lung squamous cell carcinoma collection (TCGA-LUSC), version 4 [Dataset]. The Cancer Imaging Archive (2016).

34 Aran, D., Hu, Z. & Butte, A. J. xCell: digitally portraying the tissue cellular heterogeneity landscape. Genome biology 18, 1–14 (2017).

35 Bae, S. et al. CellDART: cell type inference by domain adaptation of single-cell and spatial transcriptomic data. Nucleic acids research 50, e57–e57 (2022).

36 Bae, S. et al. STopover captures spatial colocalization and interaction in the tumor microenvironment using topological analysis in spatial transcriptomics data. bioRxiv, 2022.2011. 2016.516708 (2022).

37 Saw, P. E., Chen, J. & Song, E. Targeting CAFs to overcome anticancer therapeutic resistance. Trends in cancer 8, 527–555 (2022).

38 Zhao, L. et al. Fibroblast activation protein-based theranostics in cancer research: a state-of-the-art review. Theranostics 12, 1557 (2022).

39 Ma, C. et al. Pan-cancer spatially resolved single-cell analysis reveals the crosstalk between cancer-associated fibroblasts and tumor microenvironment. Molecular Cancer 22, 170 (2023).

40 Croizer, H. et al. Deciphering the spatial landscape and plasticity of immunosuppressive fibroblasts in breast cancer. Nature Communications 15, 2806 (2024).

41 Peng, Z., Ye, M., Ding, H., Feng, Z. & Hu, K. Spatial transcriptomics atlas reveals the crosstalk between cancer-associated fibroblasts and tumor microenvironment components in colorectal cancer. Journal of Translational Medicine 20, 302 (2022).

42 Grout, J. A. et al. Spatial positioning and matrix programs of cancer-associated fibroblasts promote T-cell exclusion in human lung tumors. Cancer Discovery 12, 2606–2625 (2022).

43 Takahashi, Y. et al. Fibrous stroma is associated with poorer prognosis in lung squamous cell carcinoma patients. Journal of Thoracic Oncology 6, 1460–1467 (2011).

44 Pellinen, T. et al. Fibroblast subsets in non-small cell lung cancer: Associations with survival, mutations, and immune features. JNCI: Journal of the National Cancer Institute 115, 71–82 (2023).

45 Bai, L., Huo, R., Fang, G., Ma, T. & Shang, Y. MMP11 is associated with the immune response and immune microenvironment in EGFR-mutant lung adenocarcinoma. Frontiers in Oncology 13, 1055122 (2023).

46 Kim, H. S., Kim, M. G., Min, K.-W., Jung, U. S. & Kim, D.-H. High MMP-11 expression associated with low CD8+ T cells decreases the survival rate in patients with breast cancer. PLoS One 16, e0252052 (2021).

47 Li, Z., Sun, C. & Qin, Z. Metabolic reprogramming of cancer-associated fibroblasts and its effect on cancer cell reprogramming. Theranostics 11, 8322 (2021).

48 Mishra, R. et al. Stromal epigenetic alterations drive metabolic and neuroendocrine prostate cancer reprogramming. The Journal of clinical investigation 128, 4472–4484 (2018).

49 Sousa, C. M. et al. Pancreatic stellate cells support tumour metabolism through autophagic alanine secretion. Nature 536, 479–483 (2016).

50 Demircioglu, F. et al. Cancer associated fibroblast FAK regulates malignant cell metabolism. Nature communications 11, 1290 (2020).

51 Comito, G., Ippolito, L., Chiarugi, P. & Cirri, P. Nutritional exchanges within tumor microenvironment: impact for cancer aggressiveness. Frontiers in oncology 10, 396 (2020).

52 Fiaschi, T. et al. Reciprocal metabolic reprogramming through lactate shuttle coordinately influences tumor-stroma interplay. Cancer research 72, 5130–5140 (2012).

53 Han, Y. et al. TISCH2: expanded datasets and new tools for single-cell transcriptome analyses of the tumor microenvironment. Nucleic acids research 51, D1425–D1431 (2023).

54 Wolf, F. A., Angerer, P. & Theis, F. J. SCANPY: large-scale single-cell gene expression data analysis. Genome biology 19, 1–5 (2018).

55 Lopez, R., Regier, J., Cole, M. B., Jordan, M. I. & Yosef, N. Deep generative modeling for single-cell transcriptomics. Nature methods 15, 1053–1058 (2018).

56 Palla, G. et al. Squidpy: a scalable framework for spatial omics analysis. Nature methods 19, 171–178 (2022).

57 Na, K. J. et al. Reciprocal change in glucose metabolism of cancer and immune cells mediated by different glucose transporters predicts immunotherapy response. Theranostics 10, 9579 (2020).

58 Fang, Z., Liu, X. & Peltz, G. GSEApy: a comprehensive package for performing gene set enrichment analysis in Python. Bioinformatics 39, btac757 (2023).

