## Supplementary Figures for "Spatial Transcriptomics Reveals Spatially Diverse Cancer-Associated Fibroblast in Lung Squamous Cell Carcinoma Linked to Tumor Progression"

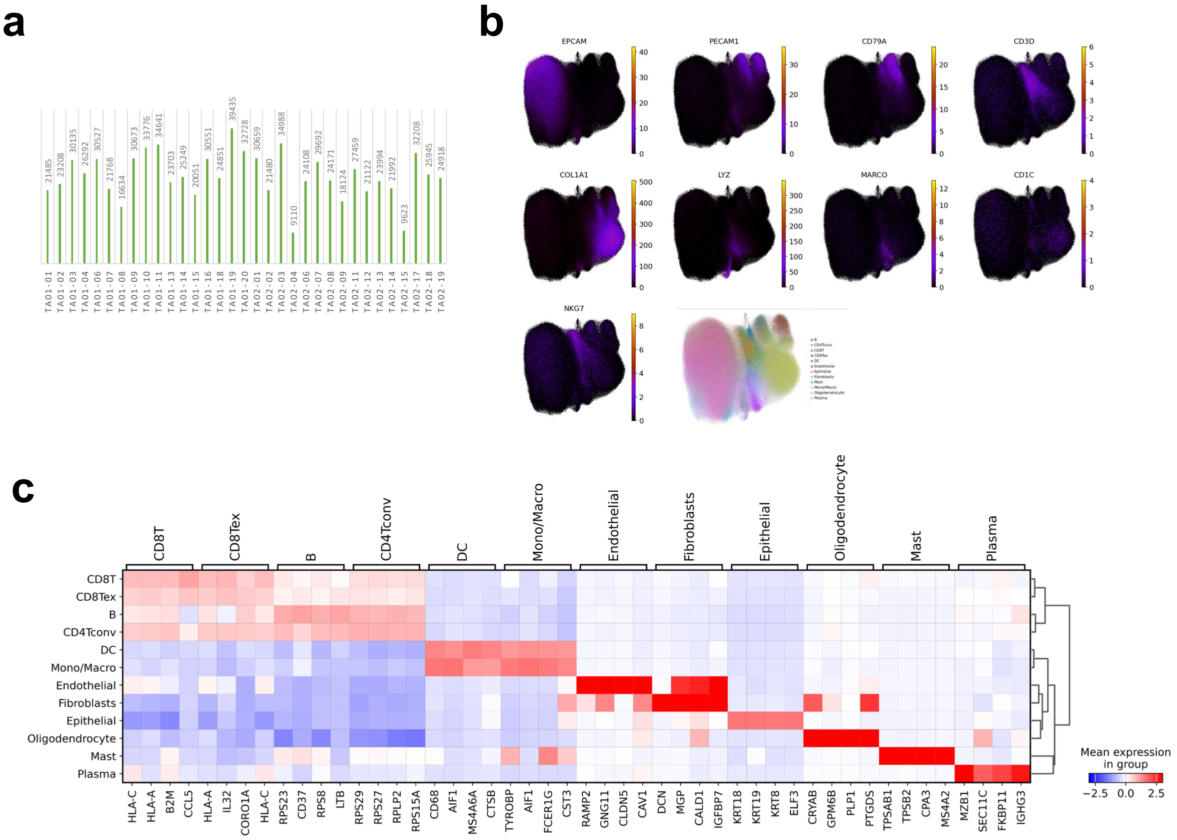

**Supplementary Figure 1. Analysis of cell type annotations in LUSC**

(a) Quantitative presentation of the total number of cells analyzed from each tissue microarray (TMA) sample. This bar plot displays the extent of cell data obtained for in-depth analysis using MERFISH.

(b) UMAP plots illustrating key markers for cell types identified in MERFISH data.

(c) Visualization of gene markers estimated from single-cell RNA sequencing data.

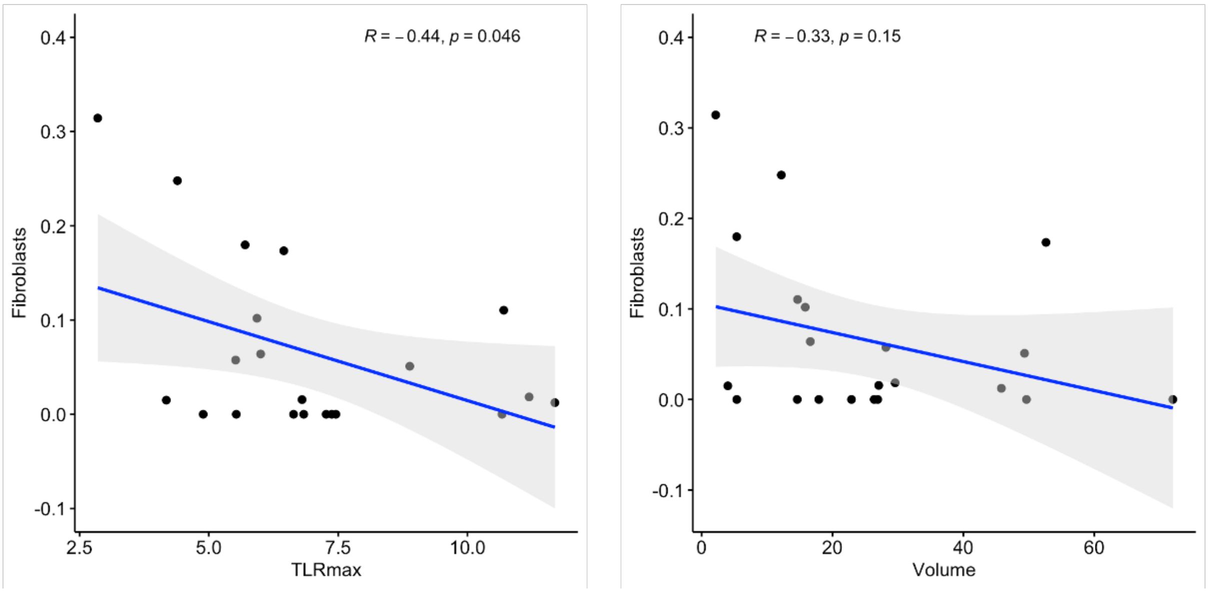


**Supplementary Figure 2. Correlation analysis between tumor volume, TLRmax and fibroblast enrichment scores from TCGA data**

The correlation analysis was performed between tumor volume and TLRmax (SUVmax normalized by liver SUVmean) against fibroblast enrichment scores calculated from bulk RNA-seq data. The tumor volue and TLRmax were obtained from the matched patients data of The Cancer Imaging Archive (TCIA). The significant negative correlation between TLRmax and fibroblast enrichment scores (rho = -0.44, p = 0.046) was, alongside a non-significant negative correlation with tumor volume (rho = -0.33, p = 0.15).

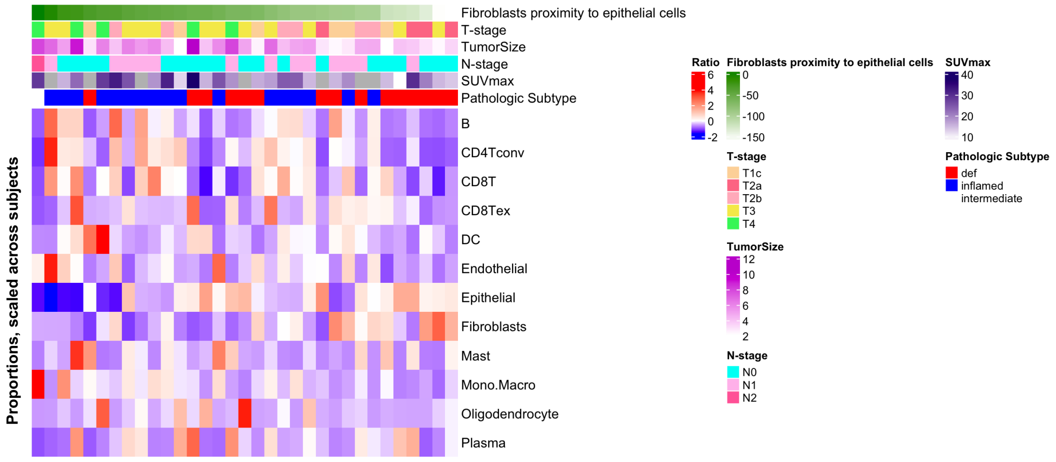


**Supplementary Figure 3. Analysis of neighborhood enrichment scores, proportions of cell types in TME and clinicopathologic features of LUSC**

A heatmap illustrates the neighborhood enrichment scores for fibroblasts and epithelial cells within the tumor microenvironment, alongside clinical variables such as tumor size, SUVmax, stage, and pathologic subtypes of LUSC. The samples are organized in descending order based on the fibroblast neighborhood enrichment scores relative to epithelial cells.

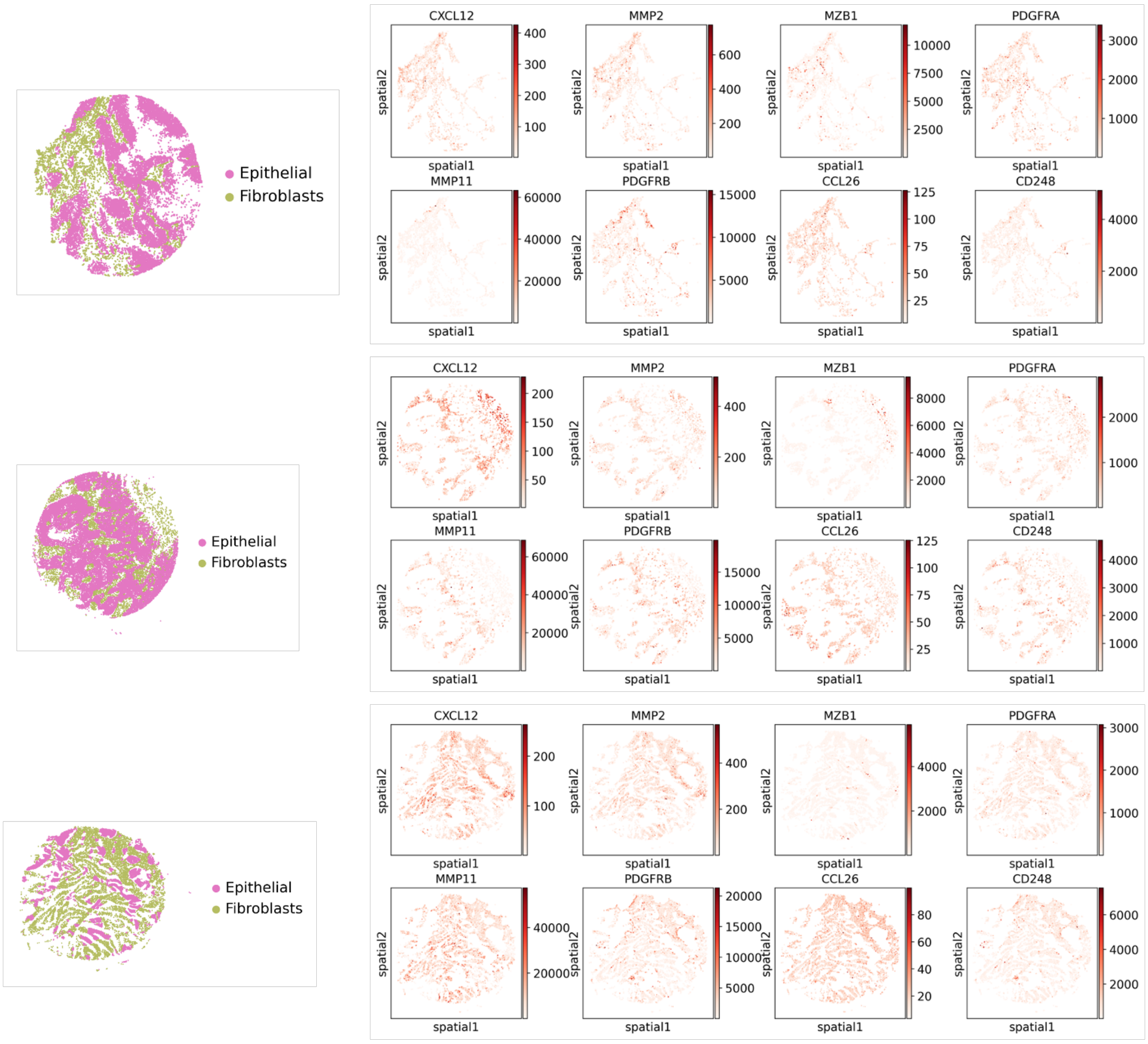


**Supplementary Figure 4. Examples of molecular markers of fibroblasts based on proximity to tumor epithelium**

As more examples, figures highlights the distinct molecular markers associated with fibroblasts categorized by their spatial proximity to tumor epithelial cells. It showcases two main groups: 'Epithelial-distant fibroblasts', characterized by markers such as CXCL12, MMP2, MZB1, and PDGFRA; and 'Epithelial-adjacent fibroblasts', identified by markers like MMP11, PDGFRB, CCL26, and CD248, which are primarily expressed at the peripheries of fibroblasts-regions.

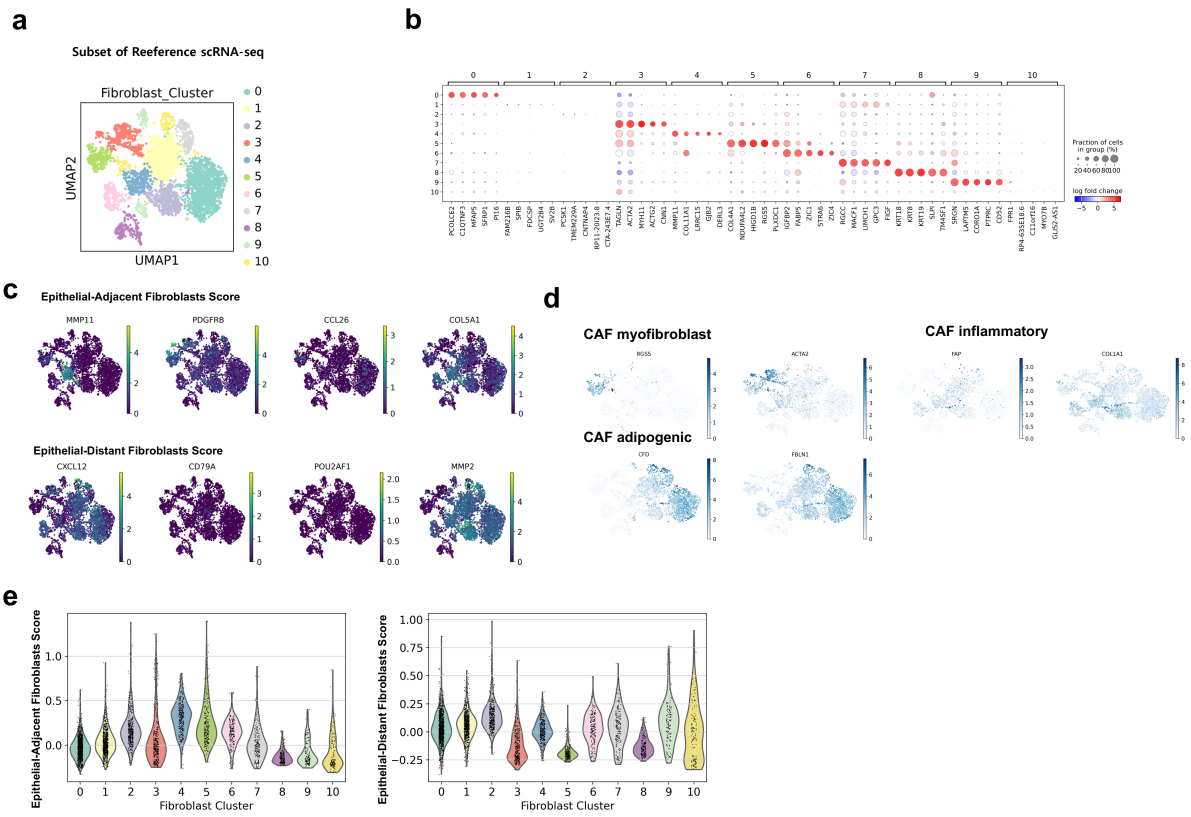


**Supplementary Figure 5. scRNA-seq profiling of fibroblasts and molecular signatures reflecting proximity to tumor epithelium**

(a) Clustering of subset scRNA-seq data of fibroblasts showed distinct groups of fibroblasts within the tumor microenvironment.

(b) According to these subclusters of fibroblasts, markers were extracted.

(c) Visualization of enrichment scores for the molecular signatures of epithelial-distant and epithelial-adjacent fibroblasts, delineating distinct clusters within the scRNA-seq dataset.

(d) Analysis of characteristic features of major cancer-associated fibroblast (CAF) states including myofibroblasts, inflammatory fibroblasts, and adipogenic fibroblasts, identified in pan-cancer CAF atlas.

(e) Specific clusters ('cluster 4' and 'cluster 5') show high scores for the epithelial-adjacent fibroblast signature, while 'cluster 0' and 'cluster 1' show high scores for the epithelial-distant fibroblast signature. These clusters exemplify the spatial and functional heterogeneity of fibroblasts within the tumor microenvironment. The primary markers defining these clusters, including IGFBP6+ CAFs in 'cluster 0' and POSTN+ myofibroblasts in 'cluster 4'.

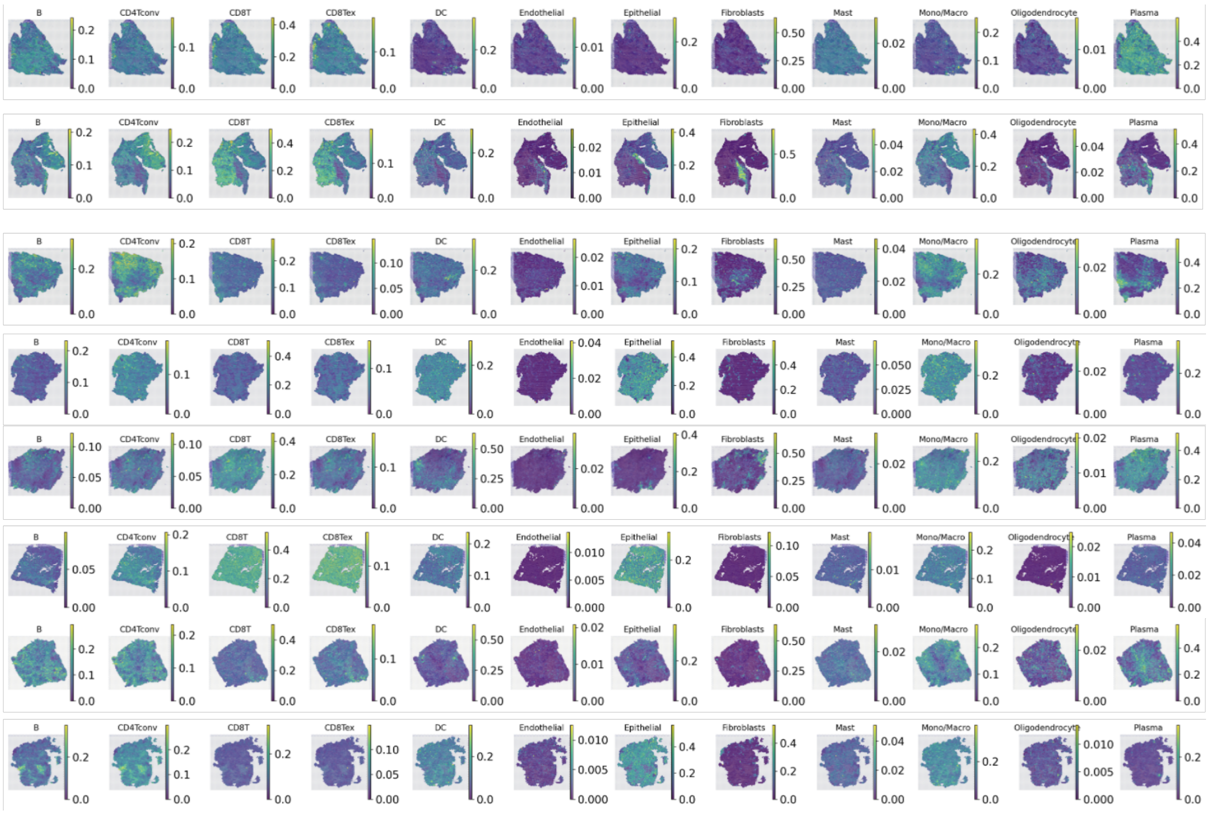


**Supplementary Figure 6. Cell type mapping in LUSC TME using Visium spatial transcriptomics.** This figure presents a spatial map of cell types within the LUSC TME, analyzed using Visium spatial transcriptomics data. The cell type distribution was characterized by the CellDART algorithm, highlighting the localization and interaction of various cell populations in the TME.
